# Endothelial-to-hematopoietic transition is induced by Notch glycosylation and upregulation of Mycn

**DOI:** 10.1101/2020.09.13.295238

**Authors:** Briane Laruy, Irene Garcia-Gonzalez, Veronica Casquero-Garcia, Rui Benedito

## Abstract

A better understanding of the molecular mechanisms driving hematopoietic stem cell (HSC) specification and expansion may enable better pharmacological strategies to produce them in sufficient numbers for transplantation. In the embryo, HSCs arise from a defined subset of arterial endothelial cells (ECs) located in the aorta–gonad–mesonephros (AGM) region that undergo endothelial-to-hematopoietic transition (EHT). Arterialization and HSC development are generally believed to require the action of Notch. Here we show that although Notch activity is initially required for arterialization, it is detrimental to subsequent EHT. Mechanistically, we show that effective EHT depends on a Mfng-induced decrease in Jag1-Notch signaling in hemogenic ECs. This causes upregulation of Mycn, an important metabolic and cell-cycle regulator that we found to be required for EHT. During the subsequent development of hematopoietic lineages, *Mycn* expression decreases and its function is taken on by the homologous *Myc* gene.

## Introduction

The adult hematopoietic system is formed by several distinct cell lineages that arise from embryonic hematopoietic stem cells (HSCs). The understanding and modulation of molecular mechanisms regulating HSC specification, expansion, and mobilization has major implications for the treatment of patients with otherwise incurable blood malignancies^1^.

The first long-term or definitive HSCs arise from hemogenic arterial ECs (HAECs) localized in the aorta-gonad-mesonephros (AGM) region of developing embryos ^2, 3, 4^. Here, a unique and yet poorly understood combination of several signalling pathways and niche factors converge to transdifferentiate a subset of dorsal aorta ECs (DAECs) to HAECs, a fraction of which give rise to rare multipotent hematopoietic progenitor cells (HPCs). This transdifferentiation process is often referred to as endothelial-to-hematopoietic transition (EHT) ^4, 5^. The biology of these hematopoietic progenitors of endothelial origin has been intensively investigated due to their unique capacity to self-renew and differentiate into all blood lineages throughout organismal lifespan or when transplanted to an organism depleted of its own blood cells ^1, 2, 3, 6^.

Genetic studies in zebrafish and mouse embryos have shown that the dorsal aorta EHT requires several transcription factors. In mice, embryos lacking Runx1, Gata2, Tal1, Gfi1, and Jag1-Notch signaling have significantly fewer aorta hematopoietic clusters ^7, 8, 9, 10, 11, 12, 13^.

Overexpression of Notch in all cells of zebrafish embryos was reported to promote formation of runx1+cmyb+ HPCs ^14, 15^. Conversely, a decrease in Notch signaling induced by morpholino-mediated knockdown of Mindbomb expression in zebrafish or deletion of the *Notch1/Rbpj* genes in mice also resulted in a significant decrease in HPC development ^9, 14, 16^. However, HPCs arise from pre-differentiated arterial ECs, and Notch signaling mutants also have significant arterial differentiation and angiogenesis defects ^17, 18, 19, 20^; therefore, these classical Notch loss-of-function studies did not address the direct role of Notch in hemogenic arterial ECs and EHT. Importantly, embryonic hematopoiesis was strongly inhibited by genetic deletion of the arterial Notch ligand Jagged1 (Jag1) ^8^, even in the absence of arterial differentiation defects. This was the first study in mice hinting at the critical requirement of Notch signalling for EHT independently of its role in the preceding arterialization process. Altogether, these results obtained in mice and zebrafish embryos indicate the existence of a conserved role of Notch signalling in promoting EHT and hematopoiesis.

In this study, we used novel multispectral conditional mosaic genetics and imaging techniques to induce differential Notch activity during EHT and fate-map its consequences *in situ*. Our work indicates that Notch activity is only required in AGM ECs in the first steps of arterialization, and that among a pool of arterial ECs with heterogenous Notch signaling levels, those with lower Notch signaling have a higher propensity to undergo EHT. By combining conditional genetic studies with single-cell RNAseq data analysis, we show that single hemogenic arterial ECs have a transient increase in the expression of the Notch glycosyltransferase Manic fringe (Mfng). Glycosylation of Notch receptors decreases the signaling ability of Jag1 ligands expressed by neighboring cells, lowering Notch signaling in signal-receiving hemogenic arterial ECs in a cell-autonomous manner. The decrease in Notch signaling leads to a strong upregulation of *Mycn* expression in arterial ECs undergoing hematopoietic transition. Genetic abrogation of Mycn function in DAECs revealed its critical requirement for EHT downstream of Notch. These Notch and Mycn functions are restricted to embryonic EHT and are not needed for the homeostasis of the adult hematopoietic system, which requires the function of the homologous *Myc* gene. Modulation of the identified molecular mechanisms may enable the generation of HSCs from ECs for therapeutic purposes.

## Results

### Notch activation in aorta ECs inhibits the formation of HPCs

Heat-shock induction of Notch activity in all cells of a zebrafish embryo has been shown to increase the expression of the hemogenic gene *runx1* and the hematopoietic gene *cmyb* ^14^. Ectopic Notch1 overexpression can also convert a venous vessel into an arterial vessel and thus a potential hematopoietic niche ^14, 17^. Work with human pluripotent stem cells (hPSCs) has shown that Notch activation during hematoendothelial differentiation potentiates EHT initiation ^21^. These results indicate that ectopic Notch activation induces HPC development to some extent; however, it is unclear whether this is a direct or indirect consequence of the role of Notch in inducing arterial differentiation. In mouse embryos it is possible to induce Notch activation specifically in ECs *in vivo*. Endothelial-specific (Tie2-Cre+) overexpression of the active Notch1 intracellular domain (N1ICD) lacking its native PEST regulatory domain, induces severe cardiovascular development and hematopoietic defects and premature death at embryonic day (E) 9.5 ^22^. Here, we used a new conditional mouse model that enables the Cre-dependent and simultaneous co-expression of fluorescent marker proteins (MbTomato and H2B-GFP) and the *Notch1* intracellular domain containing its native PEST domain (*Gt(ROSA)26Sor*^*tm1(CAG-LSL-Tomato_GFP_N1ICDP)Ben*^, *abbreviated as N1ICDP)*. This strategy results in a reportable increase in Notch signaling that more closely resembles physiological Notch signaling levels (Fig. 1a-1d). In contrast to previously published Notch gain-of-function alleles ^22, 23^, this allele, when is induced specifically in Tie2-Cre^+^ cells (*N1ICDP*^*iEC-*GOF^), produces milder angiogenesis defects, and embryos die at later stages of development (from E15.5 onward -Luo et al., under review). At E10.5, only 45% of aorta ECs in *N1ICDP*^*iEC-*GOF^ embryos were GFP^+^ (Fig. 1e, 1f), showing either that recombination of the *N1ICDP*^*iGOF*^ allele with Tie2-Cre is incomplete or that N1ICDP/GFP^+^ ECs are outcompeted during early vascular development by non-recombined cells with normal Notch signaling levels ^24^, an effect due to the negative effect of Notch overactivation on endothelial proliferation ^25, 26, 27^. These embryos, with N1ICDP overexpression in just 45% of dorsal aorta ECs, showed a 65% decrease in the average number of HPCs (CD31^+^/Kit^+^ cells) present in the dorsal aorta floor relative to their control littermates (Fig. 1g, 1h). These results indicate that sustained and higher Notch activity in mouse DAECs inhibits the formation of Kit+ hematopoietic progenitors, contrasting with previous findings using distinct experimental Notch gain-of-function approaches in zebrafish, human PSCs and mice ^14, 21, 22^.

**Figure 1.**
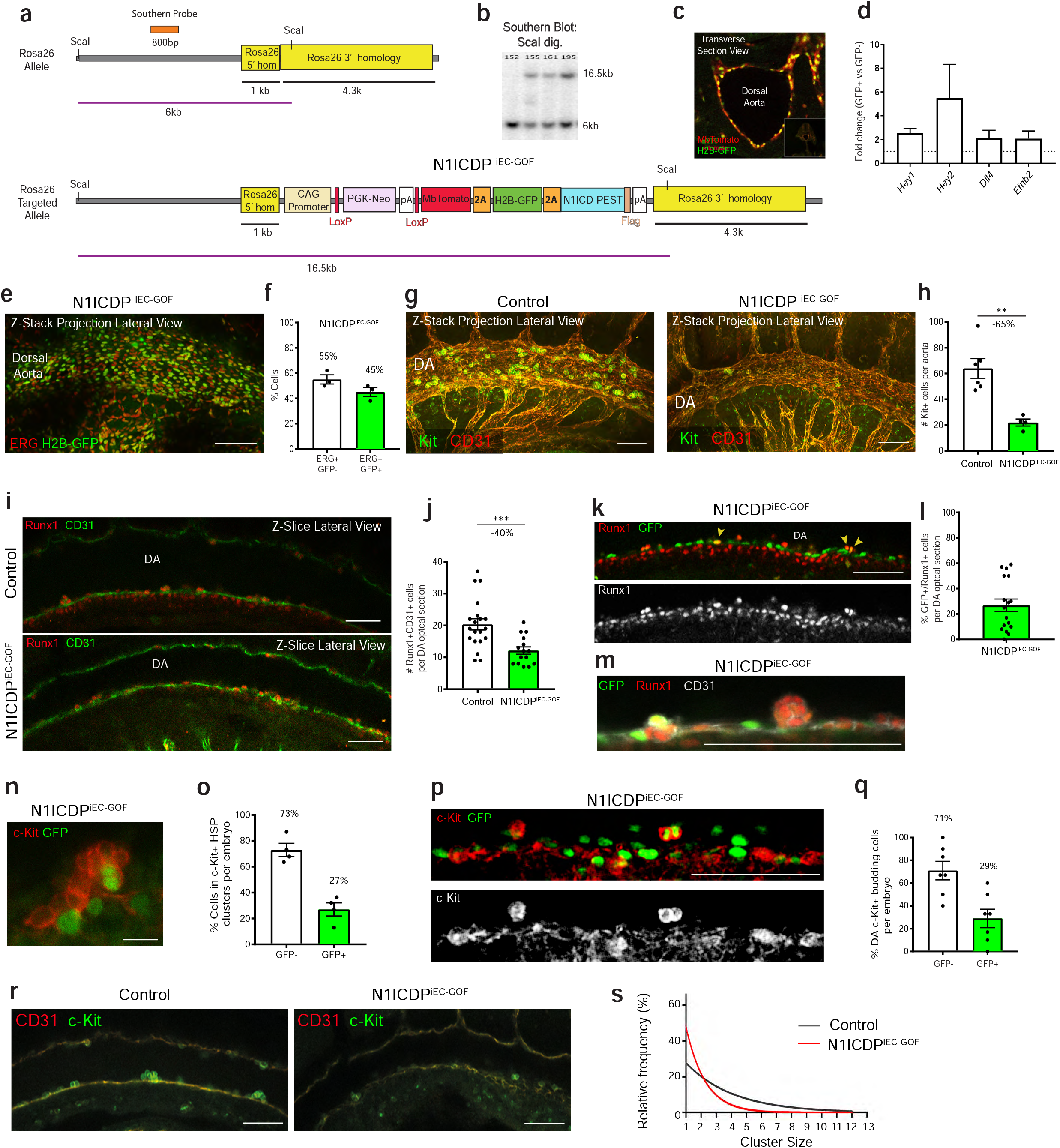
Notch activation in aorta ECs inhibits the formation of HPCs. **(a)** Genetic construct used to target the Rosa26 locus in mouse ES cells and generate mice with Cre-dependent conditional expression of MbTomato, H2B-GFP and N1ICD containing its native PEST domain. **(b)** Southern-blot showing the correct targeting of pre-selected ES cell clones (16.5kb band corresponds to the targeted allele, 6kb to the wildtype allele). **(c)** Sectional view of the DA of a *N1ICDP*^*iEC-GOF*^ mouse, showing expression of the reporter proteins MbTomato and H2B-GFP in ECs. **(d)** qRT-PCR analysis showing increased expression of the Notch target genes Hey1and Hey2 and the arterial markers Dll4 and Efnb2 in *N1ICDP*^*iEC-GOF*^ ECs (n=3 pooled litters, sorted for CD31-APC+/MbTomato+GFP+ cells and CD31+/MbTomato-GFP-cells). (**e and f**) Z-stack projection of confocal scans of the immunostained DA, showing that only a fraction of ECs (ERG+) express the *N1ICDP*^*iEC-GOF*^ allele (GFP+) (n=3 mutant embryos). (**g and h**) Z-stack projection of confocal scans of the immunostained AGM showing a decrease in the number of Kit+ cells associated with DAECs (CD31+) (n=6 sibling control and n=4 mutant embryos). (**i and j**) Confocal optical sections of immunostained DAs showing a reduction in the frequency of Runx1+/CD31+ cells in *N1ICDP*^*iEC-GOF*^ DA (n=4 sibling control and n=3 mutant embryos). (**k and l**) Confocal optical section of the *N1ICDP*^*iEC-GOF*^ DA floor, showing that only a small fraction of N1ICDP+/GFP+ DAECs are also Runx1+ (n=4 mutant embryos). (**m**) Confocal images of immunostained aortas showing that N1ICDP+/GFP+ DAECs do not form large Runx1+ clusters. (**n and o**) High magnification confocal image showing that most Kit+ cells forming DA hematopoietic clusters are GFP-(n=4 mutant embryos). (**p and q**) Confocal images of the immunostained DA floor showing that only a relatively small fraction of hemogenic (Kit+) budding ECs are GFP+ (n=7 mutant embryos). (**r and s**) Confocal optical sections of immunostained DA showing a reduction in the frequency of Kit+/CD31+ cells in *N1ICDP*^*iEC-GOF*^ DA (n=4 sibling control and n=4 mutant embryos). Scale bars, 100 μm except for n, 20 μm. Error bars indicate SEM. **p < 0.01, ***p < 0.001.

To obtain a higher resolution view of the impact of increased dorsal aorta Notch activity on HPC formation, we first counted the number of CD31^+^ Runx1^+^ cells present in the AGM dorsal aorta floor of *N1ICDP*^*iEC-*GOF^ embryos. Runx1 is one of the most important markers of HAECs and is essential for EHT. At E10.5, embryos with increased Notch activity in DAECs had 40% fewer CD31^+^/Runx1^+^ cells (average of 12 cells per section) than their control littermates (average of 20 cells per section) (Fig. 1i, 1j). Only 30% of endothelial *N1ICDP*^*iEC-*GOF^/GFP^+^ cells were also Runx1^+^ (Fig. 1k, 1l), showing that an increase in dorsal aorta Notch signaling inhibits but does not abolish the initial endothelial commitment to the hematopoietic lineage (Runx1^+^). However, *N1ICDP*^*iEC-*GOF^ /GFP^+^/Runx1^+^ cells did not form large hematopoietic clusters (Fig. 1m).

We next compared the frequencies of control (GFP^−^) and mutant (GFP^+^) *N1ICDP*^*iEC-*GOF^ cells, in the endothelial (ERG^+^) and HPCs (Kit^+^) populations. Whereas 45% of DAECs were N1ICD-GFP^+^ (Fig. 1e, 1f), only 27% of cells in Kit^+^ aortic hematopoietic clusters were GFP^+^ (Fig. 1n, 1o). This difference suggests either that DAECs with higher Notch signaling are unable to complete EHT or that they proliferate less after they undergo EHT and become Kit^+^ HPCs. To distinguish the effect of Notch on these consecutive events, we first counted GFP^+^ positive cells in the population of single budding endothelial Kit^+^ cells, which are the putative founder cells undergoing EHT that later form the HPC clusters. Whereas 45% of DAECs were N1ICD-GFP^+^, only 29% of budding Kit+ DAECs were *N1ICDP*^*iEC*^/GFP^+^ (Fig. 1p, 1q), suggesting once again that sustained Notch activation in single dorsal aorta ECs impairs, but does not abolish, Runx1 expression and hemogenic endothelial budding. To determine if sustained Notch activation also impacts the expansion of HPC clusters after the budding event, we quantified the size (number of Kit^+^ cells per cluster) of all Kit^+^CD31^+^ clusters in control and *N1ICDP*^*EC-iGOF*^ aortas. The results show that increased Notch activity significantly reduced the frequency of larger HPC clusters (Fig. 1r, 1s). The sum of these data indicates that even though Notch activation in dorsal aorta ECs inhibits the initial steps of conversion to HAECs (acquisition of *Runx1* expression and cell budding), its sustained activity particularly affects the early expansion of already committed DA Kit^+^ hematopoietic progenitor cell clusters.

### Dorsal aorta ECs with lower Notch signalling more often undergo EHT

Several genetic studies have shown that the full loss of Dll4–Notch1–Rbpj signaling severely affects the development of the dorsal aorta ^19, 28, 29, 30^ and the subsequent EHT ^9, 16^. We therefore sought to avoid the problem of lack of arteriogenesis in global or EC-specific Notch mutants by instead analyzing embryos expressing an inducible fluorescent genetic Notch mosaic allele (*ifg-Notch-Mosaic*)^31^. These Notch mosaic embryos develop normally due to the relatively low frequency of mutant cells having a sustained decrease or increase in Notch signalling ^24, 31^. Embryos carrying the *iChr2-Notch-Mosaic* and *Tie2-Cre* alleles (abbreviated here as *Notch-Mosaic*^*iEC*^) will produce (starting at E7.5-E8.0) a mosaic of ECs with normal (Cherry+), low (H2B-GFP+ and DN-Maml1+), or high (HA-H2B-Cerulean+ and N1ICD-PEST+) Notch activity (Fig. 2a). We previously showed that ECs of *Notch-Mosaic*^*iEC*^ embryos expressing different fluorescent proteins, and hence having distinct Notch signaling levels, have distinct proliferative and sprouting abilities ^24^. Importantly, cells expressing H2B-GFP and DN-Maml1 have more Notch signalling than cells with full loss of *Dll4* or *Rbpj* and are able to form arteries ^24^. We also previously showed that the baseline or initial recombination/expression frequency of the *iChr2-Notch-Mosaic* allele after Cre plasmid transfection *in vitro* is 44% Cherry+, 32% GFP+ and 24% Cerulean+ ^31^. Here, we found that the dorsal aorta of E9.5 *Notch-Mosaic*^*iEC*^ embryos was formed by 65% Cherry+, 21% GFP+ and 14% Cerulean+ cells (Fig. 2b, 2c), likely reflecting the lower tendency of cells with low Notch (DN-Maml1^+^/GFP^+^) to become incorporated into arteries and the lower proliferation of ECs with high Notch signalling (N1ICDP/Cerulean^+^) ^24, 32^. Between E9.5 and E10.5 The frequency of the Cherry^+^ EC population increased to 74%, while the GFP^+^ and Cerulean^+^ frequencies decreased to 16% and 10%, respectively (compare Fig. 2c with 2e, EC bars). Analysis of the mosaic frequency in the Kit^+^ HPC population attached to the dorsal aorta revealed that the DN-Maml1/GFP^+^ population was 68% more frequent than in ECs (27% of E10.5 Kit+ HPCs versus 16% of E10.5 DAECs), whereas the N1ICDP/Cerulean^+^ population was 58% less frequent (4% of Kit+ HPCs versus 10% of E10.5 DAECs) (Fig. 2e). These results suggest that dorsal aorta ECs with lower Notch activity have an increased propensity to undergo endothelial-to-hematopoietic transition and become Kit+ HPCs, whereas cells with higher Notch signaling tend not to form HPCs.

**Figure 2.**
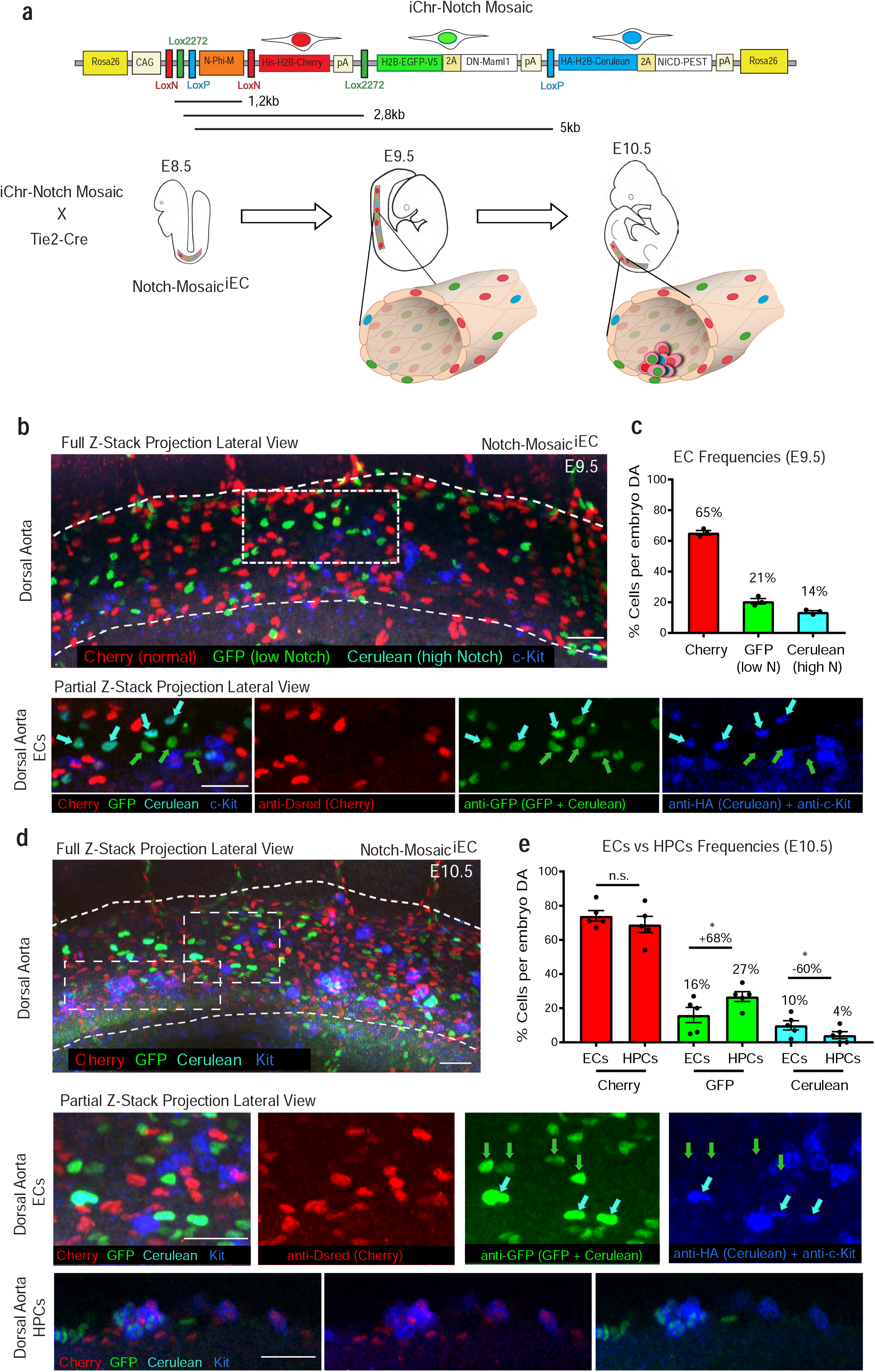
Analysis of Notch multispectral genetic mosaics. **(a)** *iChr2-Notch-Mosaic* DNA construct with the genetic distance (kb) between different LoxP sites indicated. In embryos carrying the *iChr2-Notch-Mosaic* and *Tie2-Cre* alleles a genetic mosaic is induced at E8.0-E8.5 in ECs of developing DAs. Due to the mutually exclusive recombination nature of the distinct loxP sites in the construct, different cells will be labeled with one of the three possible chromatin (H2B) labels. H2B-Cherry+ cells will be controls, H2B-GFP+ cells will co-express DN-Maml1 which lowers Notch signaling; and HA-H2B-Cerulean+ cells will co-express N1ICD-PEST which activates Notch signaling. **(b)** Z-stack projection of confocal scans of the *Notch-Mosaic*^*iEC*^ DA at E9.5, immunostained with anti-Dsred (detects H2B-cherry), anti-GFP (detects H2B-GFP and HA-H2B-Cerulean), anti-HA (detects only HA-H2B-Cerulean) and anti-Kit. The combination of the 4 signals allows the distinction of all cell types, as can be seen in the high magnification views of the boxed area. **(c)** Quantification of the distinct cell populations in mouse DAs (n=3 embryos). **(d)** Z-stack projection of confocal scans of *Notch-Mosaic*^*iEC*^ DA at E10.5. Boxed areas show the different endothelial and HPC (Kit+) populations. **(e)** Quantification of the distinct cell populations in mouse DAs (n=5 embryos). ECs with low Notch (GFP+) tend to form more HPCs, whereas ECs with high Notch (HA+) rarely form Kit+ clusters. Scale bars, 45 μm. Error bars indicate SEM. *p < 0.1.

### Endothelial loss of Jag1 phenocopies the N1ICDP overexpression effect in EHT

Unlike other Notch signaling regulators (Notch1, Rbpj and Dll4), Jag1 function is dispensable for the development of the dorsal aorta ^33^. Global loss of Jag1 does not affect arteriogenesis but results in a specific impairment of Gata2 expression and AGM hematopoiesis, a phenotype that was attributed to the loss of Jag1-Notch activation in DAECs, even though no significant Notch activity differences were reported in these mutants ^8^. To assess the phenotype of Jag1 loss-of-function specifically in aorta ECs at high resolution, we generated compound Tie2-Cre Jag1^fl/fl^ embryos (abbreviated *Jag1*^*iEC-KO*^) and counted the number of Kit^+^ cells in the DA at E10.5. *Jag1*^*iEC-KO*^ mutants had 69% fewer Kit^+^ HPCs in the DA than their control littermates (Fig. 3a, 3b). These results confirm that EHT requires Jag1 function in ECs. However, the similarity of the phenotype of these mutants to that of the *N1ICDP*^*iEC*^ embryos raises the question of whether Jag1, a canonical Notch ligand, could act as an inhibitor of Notch signaling in hemogenic aorta ECs as was previously shown to occur during angiogenesis ^25^. It was previously shown that the transcriptional expression of several canonical Notch targets is upregulated in CD31+Kit- and CD31+Kit+ cell explants derived from E10.5 embryos with global Jag1 deletion ^34^. Since *Dll4* expression is also induced after Notch activation in ECs ^25, 35^, we stained with a Dll4 antibody to detect any differences in Notch activation levels in the DA of these embryos. Dll4 immunosignals, particularly endocytosed puncta induced by its engagement with Notch receptors, were significantly increased in *Jag1*^*EC-KO*^ mutant aorta endothelium (Fig. 3c, 3d). qRT-PCR analysis of *Jag1*^*EC-KO*^ mutant ECs confirmed increased expression of canonical endothelial Notch target genes (Fig. 3e). These data confirms that loss of endothelial Jag1 increases Dll4-Notch activation in the dorsal aorta endothelium, which inhibits the formation of HPCs, as in *N1ICDP*^*iEC*^ mutants.

**Figure 3.**
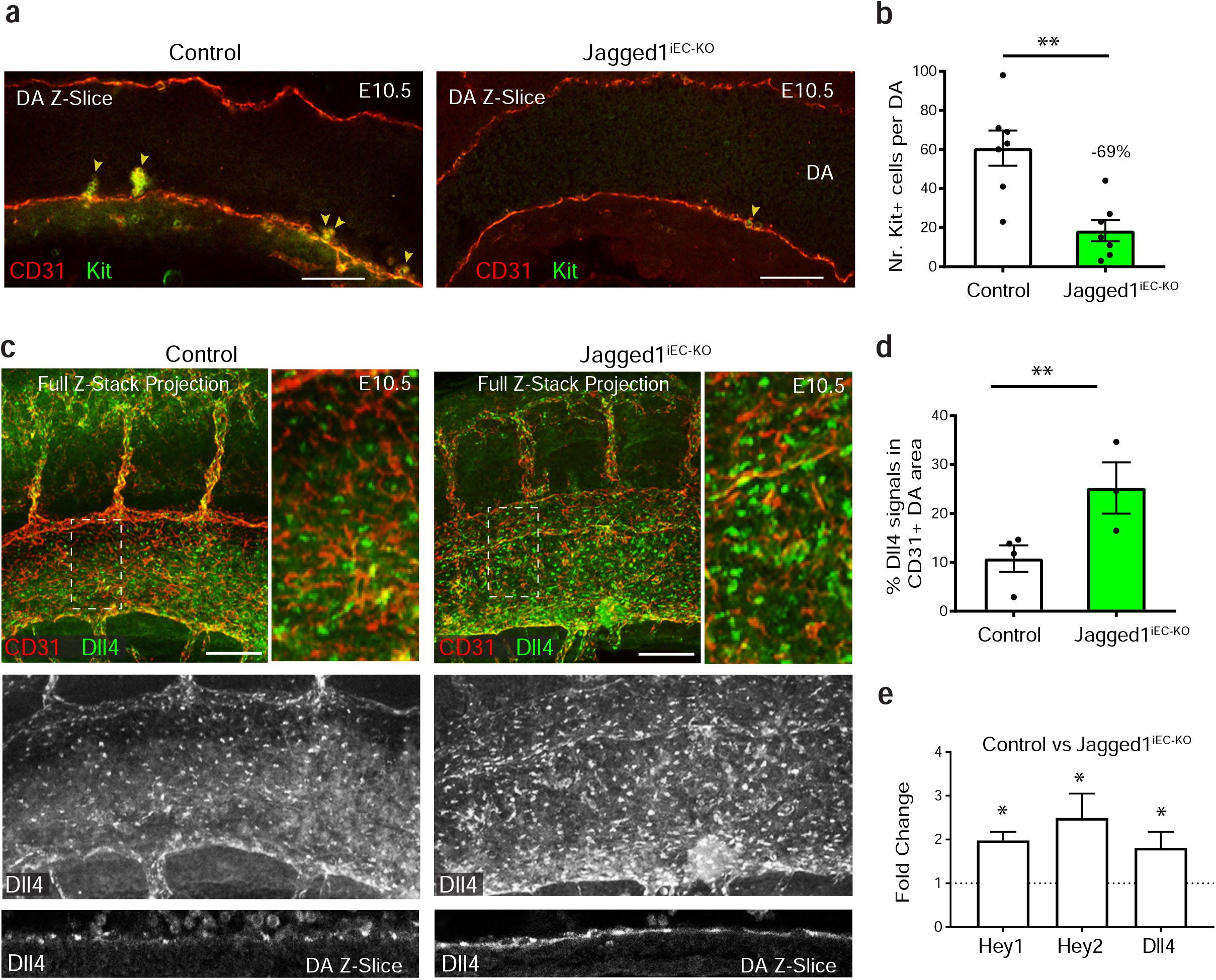
Jag1 inhibits EHT by downregulating Notch signaling. **(a)** Z-stack projection of confocal scans of control and *Jag1*^*iEC-KO*^ DAs at E10.5, immunostained for CD31 and Kit. **(b)** Chart showing that the loss of Jag1 ligand in DAECs induces a decrease in the number of Kit+ cells in the DA floor (n=7 sibling control and mutant embryos). **(c)** Full or partial Z-stack projection of control and *Jag1*^*iEC-KO*^ DAs at E10.5, immunostained for CD31 and Dll4. **(d)** Chart showing that the loss of Jag1 ligand in DAECs increases the frequency of Dll4 immunosignals in the DA (n=4 sibling control and n=3 mutant embryos). **(e)** qRT-PCR showing an increase in the expression of canonical Notch target genes in AGM ECs after deletion of Jag1. Scale bars, 100 μm. Error bars indicate SEM. *p < 0.1, **p < 0.01.

### Mfng expression in HAECs induces EHT by decreasing Jag1-Notch signalling

The results presented so far raise the question of why Jag1 is important for the regulation of Notch activity in HAECs located in the AGM but not for the regulation of Notch signaling during arterialization, since *Jag1*^*EC-KO*^ mutants have normal arteries. The ligand Dll4 is strongly expressed by angiogenic and dorsal aorta ECs and is a strong regulator of Notch signaling during capillary angiogenesis and arterialization ^19, 20^. Dll4 signalling function cannot be compensated by Jag1 in these settings, suggesting that Jag1 may be less expressed or less potent than Dll4 in activating Notch signalling.

Dll4 and Jag1 are expressed in the AGM aorta endothelium beneath the Kit+ hematopoietic clusters; however, in contrast with arteriogenesis, loss of Jag1 cannot be compensated by Dll4 in these cells during EHT ^8, 34^. This prompted us to question whether the AGM hemogenic endothelium (HE) specifically expresses a modifier that increases the importance of Jag1 for Notch signaling in these cells. The Fringe family of glycosyltransferases (Lfng, Mfng, and Rfng) are broadly expressed and are able to modify the signaling strength of Notch ligands ^36, 37, 38^. By glycosylating the Notch receptors they inhibit the signaling ability of Jagged ligands while promoting the signaling ability of Delta ligands ^25, 36, 37, 38, 39^.

To assess the expression of Notch signalling components and its modifiers (Fringe genes) during EHT, we analyzed a recently published single-cell RNA-sequencing dataset of mouse aorta cell types ^40^. This dataset (series record GSE112642) includes the transcriptome of aorta ECs, hemogenic ECs, cells undergoing EHT, intra-aortic Kit+ hematopoietic clusters and more defined hematopoietic cells. Our bioinformatic analysis was able to produce a similar clustering identification with the exception that we found a total of 9 cell clusters (Fig. 4a, 4b). Clusters C0 to C6 were obtained from mouse AGM, whereas clusters 7 and 8 were obtained from yolk sac ^40^. C0 is formed by Cdh5+Gfi1-Kit-ECs, C1 and C2 are mostly composed of Cdh5+Gfi1+Kit-hemogenic ECs. C3 and C4 are composed of cells undergoing EHT (Cdh5+Gfi1+Kit+). C5 is composed of intra-aortic hematopoietic clusters (CD31+Gpr56+Kit+); and C6 to C8 represent more committed Kit+ hematopoietic progenitor cells obtained from the yolk sac (Fig. 4a-4c). Pseudotime analysis of these clusters revealed high expression of the arterial endothelial markers *Cdh5, Gja5* and *Sox17* in C0 (DAECs), with expression then gradually decreasing as cells progress toward a hematopoietic fate. Conversely, expression of the hematopoietic genes Rac2 and Myb becomes notable after C5 and gradually increases until C7 and C8, which are mainly formed by cells of a more defined hematopoietic lineage (Fig. 4c, 4d). The HE marker Gfi1 ^40, 41, 42, 43^, is highly expressed in clusters C1 and C2 (Fig. 4e, 4f). C3 and C4, formed by cells undergoing EHT, highly express Ikzf2, Kit, Ogdh and Runx1 (Fig. 4e, 4f).

**Figure 4.**
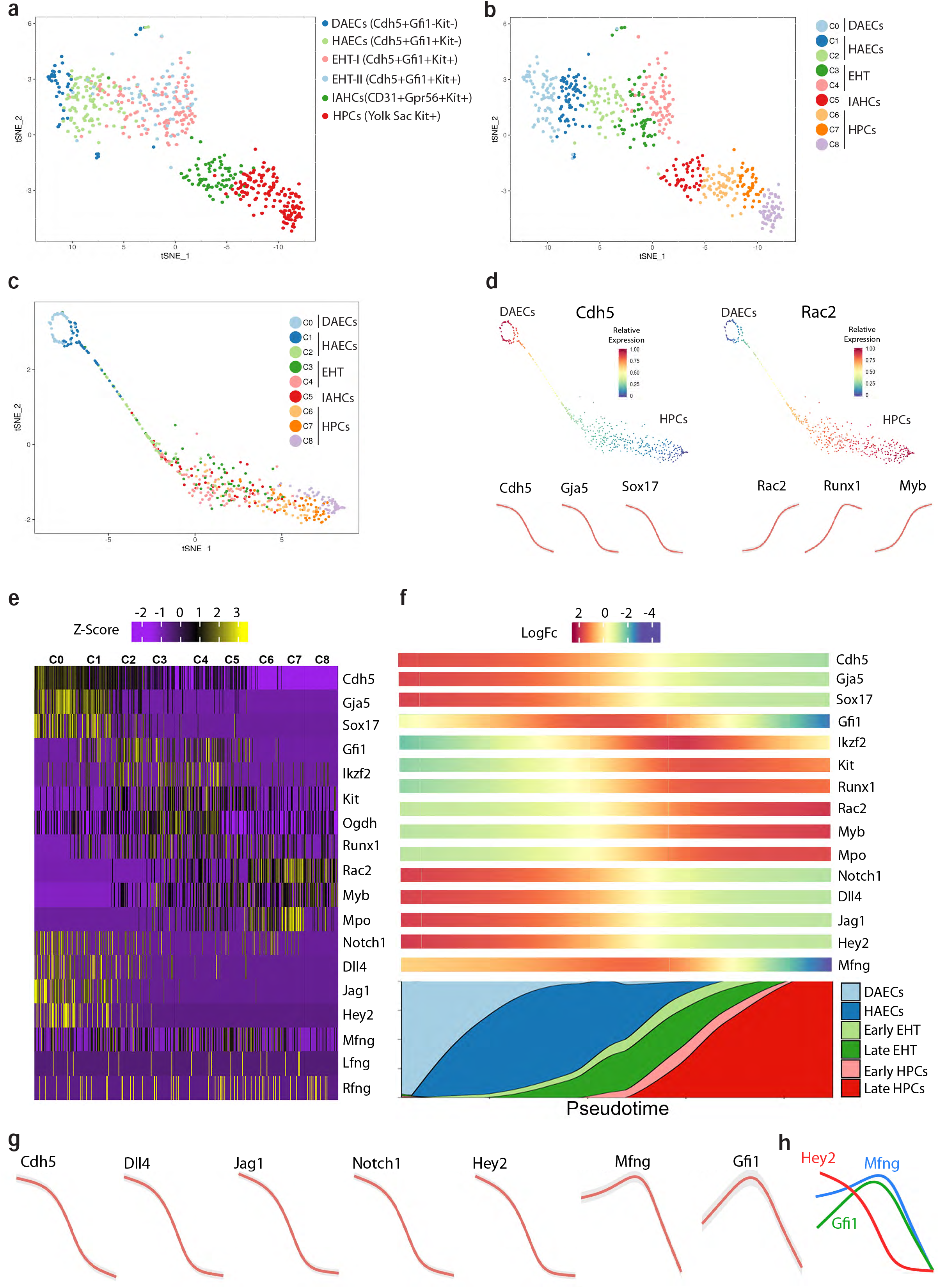
scRNAseq data analysis identifies Mfng upregulation in the HE. (a) tSNE plot of the different cell sample groups extracted by Baron et al. Color indicates the distinct groups of cells isolated from E10-E10.5 embryos or yolk sacs with the indicated markers. (b) tSNE plot showing the different clusters of cells identified bioinformatically. (c) tSNE alignment of cells along a differentiation trajectory according to the Palantir algorithm. (d) Variation of individual gene expression across pseudotime; *Cdh5, Gja5*, and *Sox17* are more highly expressed in single ECs, whereas *Rac2, Runx1*, and *Myb* are more highly expressed in single HPCs. (e) Relative expression of the indicated genes in single cells within the 9 identified clusters. (f) Left-to-right pseudotime alignment of selected genes expression within the identified single cell clusters (C0 to C8). (g, h) Relative gene expression level curves across pseudotime for selected endothelial (Cdh5), Notch modulators (Dll4, Jag1, Notch1, Hey2, Mfng) and hemogenic endothelium (Gfi1) genes.

This comparative pseudotime analysis shows that Notch pathway genes are strongly expressed in defined arterial ECs (C0), but that their expression decreases in the HE (C1 and C2) and is very low in cells undergoing EHT and in hematopoietic lineages. Analysis of Fringe genes showed that *Mfng* (but not *Lfng* or *Rfng*) has a similar expression profile to *Gfi1*, with high expression in C1 and C2, clusters formed by HAECs (Fig. 4f-4g). The scRNAseq dataset also shows that *Mfng* expression increases in Gfi1+ HAECs coinciding with the significant decrease in the expression of the Notch target gene *Hey2* (Fig. 4h). This single cell expression analysis identifies Mfng as a candidate modifier of ligand-Notch signaling in HE cells.

To investigate the impact of Mfng on EHT, we elevated endothelial Mfng expression by crossing *Tie2-Cre* mice with mice expressing Cre-inducible *Rosa26-Lox-Stop-Lox-Mfng* (abbreviated here as *Manic Fringe*^*iEC-GOF*^). Analysis at E10.5 showed that *Manic Fringe*^*iEC-GOF*^ embryos had more Kit^+^ cells in the DA than their control littermates (Fig. 5a, 5b). This suggests that the outcome of Mfng expression in the DA is Notch signaling inhibition, which we show above enhances EHT (Fig. 2e). Supporting this conclusion, immunostaining of E10.5 DA for Dll4 revealed reduced expression and frequency of bright puncta corresponding to endocytosed ligand-receptor pairs (Fig. 5c, 5d), which are known to be induced by Notch activity in ECs ^25, 44^. qRT-PCR analysis of FACS-sorted ECs from Mfng gain-of-function embryos confirmed the decreased expression of the canonical Notch targets Hey1 and Hey2 (Fig. 5e). These data are compatible with a model in which Mfng expressed in the HAECs glycosylates Notch receptors, inhibiting signaling from Jag1 ligands expressed on neighboring cells. However, in the aorta, ECs express both Jag1 and Dll4 ligands, and Mfng expression should also promote Dll4 signaling, while inhibiting Jag1 signaling ^36^. Interestingly, we found that AGM DAECs express significantly more Jag1 than Dll4 (Fig. 5f and 5g), and these ligands are able to compete with each other for Notch binding and signaling. We showed previously that in the presence of Mfng-glycosylated Notch receptors, Jag1 ligands have weaker signaling activity and sequester Notch receptors, inhibiting the binding and function of the stronger Dll4 ligands^25^. The relatively high expression of the weak Jag1 ligands in the AGM DA thus explain why Mfng overexpression decreases overall Notch signaling in this particular endothelium and why Jag1 is important in EHT and not in arterialization.

**Figure 5.**
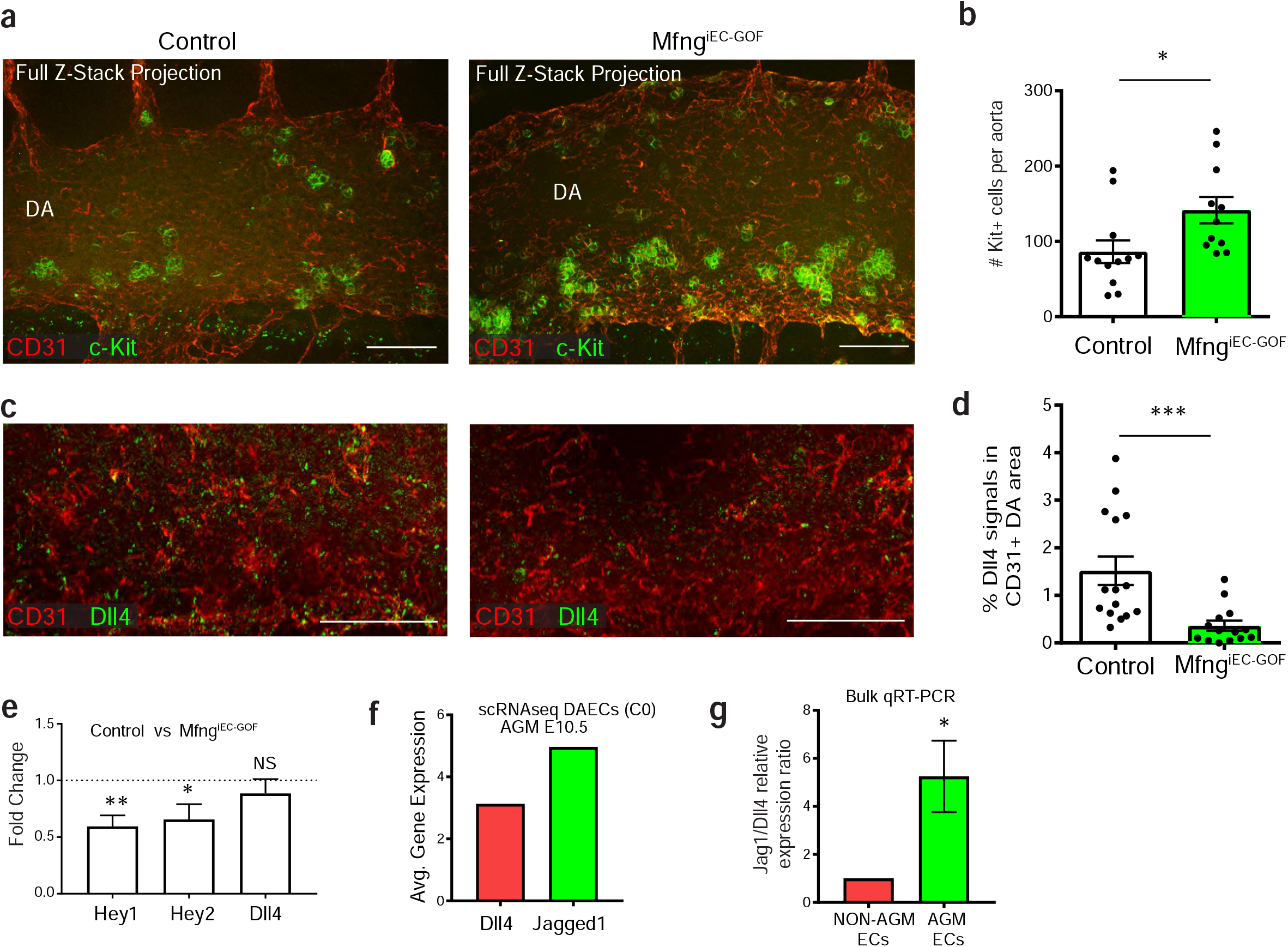
Mfng induces EHT by negatively modulating Jag1-Notch signaling. a) Z-stack projection of confocal scans of control and *Mfng*^*iEC-GOF*^ DAs at E10.5, immunostained for CD31 and Kit. b) Mfng gain-of-function induces an increase in the number of Kit+ aortic hematopoietic clusters (n=6 sibling control and mutant embryos). c) Z-stack projection of confocal scans of control and *Mfng*^*iEC-GOF*^ DAs at E10.5, immunostained for CD31 and Dll4. Note that strong puncta correspond to internalized Dll4 ligand. d) Mfng gain-of-function induces a decrease in the number of immunosignals for Dll4 (n=4 sibling control and mutant embryos). e) qRT-PCR analysis of Notch genes in AGM ECs isolated from control and *Mfng*^*iEC-GOF*^ embryos (n=3 sibling control and mutant embryos). f) Relative expression of Jag1 and Dll4 mRNA in single AGM ECs (cluster C0) isolated from embryos^40^. g) Relative expression of Jag1 and Dll4 in AGM and non-AGM ECs collected from wildtype embryos. AGM ECs express more Jag1 than Dll4, relative to NON-AGM ECs (samples collected from 3 independent pools of embryos). Scale bars, 100 μm. Error bars indicate SEM. NS, non-significant; *p < 0.05; **p < 0.01; ***p < 0.001.

### Aorta ECs with low Notch signaling upregulate *Mycn* during EHT

A distinguishing feature of DAECs and cells undergoing EHT is proliferation. Most DAECs at E10.5 are quiescent. However Kit+ cells in the DA are highly proliferative (Fig. 6a), as also shown before ^45, 46^. scRNAseq data analysis of genes normally enriched in cycling cells confirmed that cells undergoing EHT are more proliferative than most AGM DAECs (Fig. 6b, 6c). The loss of Notch signaling in non-cycling or quiescent ECs is also associated with an increase in proliferation ^24, 32, 47^. We therefore reasoned that lower Notch signaling in the DA may facilitate EHT in part by promoting cell-cycle entry. We recently found that Dll4–Notch signaling negatively regulates the expression of *Myc* and several of its target genes in coronary ECs (Luo et al., under review), whereas Notch has been shown to positively regulate proliferation and *Myc* in hematopoietic cells ^48^. Therefore, we analyzed if the expression of *Myc* was also upregulated in Cx40+ DAECs with lower Notch signaling (DN-Maml1/GFP^+^) during EHT. *Myc* expression was not altered significantly in Cx40+/DN-Maml1+/GFP^+^ ECs (Fig. 6d, 6e); however, analysis of the scRNAseq data revealed that *Mycn*, but not *Myc*, was among the top 20 genes enriched in the EHT clusters (Fig. 6f, 6g). *Mycn* is a closely related homolog of *Myc*, and can subsitute *Myc* function when expressed in the same cells ^49^. Comparative analysis of *Mycn* expression in DAECs with normal (Cx40+/Cherry+) and lower Notch signaling (Cx40+/DN-Maml1+/GFP^+^) revealed a 6-fold increase in the latter (Fig. 6e). These results led us to hypothesize that *Mycn* upregulation induced by the low Jagged1–Notch signaling activity in Mfng-expressing hemogenic DAECs could be a key driver of EHT. Supporting this idea, pseudotime analysis of the published scRNAseq data showed that the Notch target gene *Hey2* is downregulated in Mfng-expressing hemogenic ECs and that this is followed by upregulation of Mycn (Fig. 6h).

**Figure 6.**
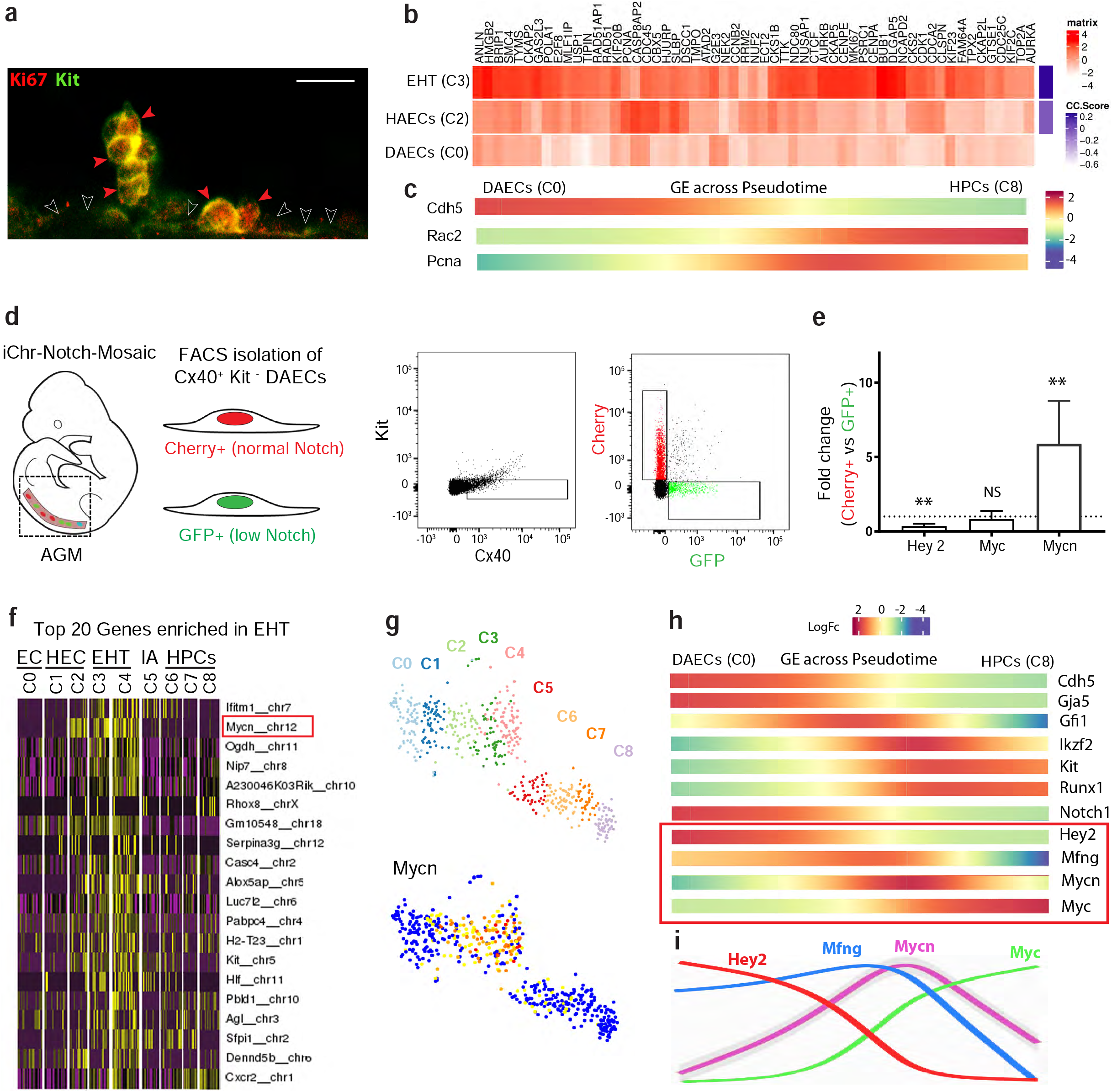
Increased expression of proliferation marker genes and Mycn during EHT. (a) Confocal micrograph showing Ki67 and Kit immunosignals in hematopoietic clusters adjacent to aorta ECs. (b) Heatmap showing the relative expression of cell-cycle genes in single cell clusters formed by DAECs (C0), HAECs (C2) and cells undergoing EHT (C3). HAECs and cells undergoing EHT are more proliferative. (c) Pseudotime alignment showing that expression of the cell-cycle marker Pcna is higher during EHT and relatively lower in ECs (Cdh5+) and HPCs (Rac2+). (d) Isolation of Cx40+Kit-DAECs with normal (Cherry+) and lowered (GFP+) Notch signaling from E10.5 embryos AGM. (e) qRT-PCR analysis showing lower expression of Hey2 and higher expression of Mycn in DN-Maml1+ DAECs (collected from 2 independent pools of embryos containing 6 or 8 embryos each). (f) Cluster analysis of the top 20 genes enriched in single cells undergoing EHT (C2, C3 and C4), showing that Mycn is the second most differentially expressed gene. (g) t-SNE plot showing the different clusters (as in Fig. 4b) and the higher expression of Mycn in single cells within C2-C4 (EHT). (h) Single cell gene expression (GE) across pseudotime showing the higher expression of Mycn after Mfng upregulation and Hey2 downregulation. Myc expression only increases in cells with hematopoietic characteristics. Scale bar, 20 μm. Error bars indicate StDev. NS, non-significant; **p < 0.01.

### Mycn is a key regulator of EHT

Existing published evidence points towards a role of Myc, not Mycn, in EHT and hematopoiesis. Dubois and colleagues showed that embryos lacking the Myc function in endothelial and hematopoietic cells (Tie2-Cre+) had a 76% decrease in CD45+ blood cells at E11, but a relatively high fraction of hematopoietic progenitors ^50^, whereas He and colleagues, using a similar mouse model, found no hematopoietic clusters in sections of the aorta using H&E staining^51^. Given these contradicting results, we decided to further investigate the role of Myc in the EHT process using our 3D whole aorta imaging technique. In *Myc Tie2-Cre (Myc*^*iEC-KO*^) E10.5 embryos *w*e observed no differences in the number of Kit^+^ HPCs present in the DA floor (Fig. 7a, 7b).

**Figure 7.**
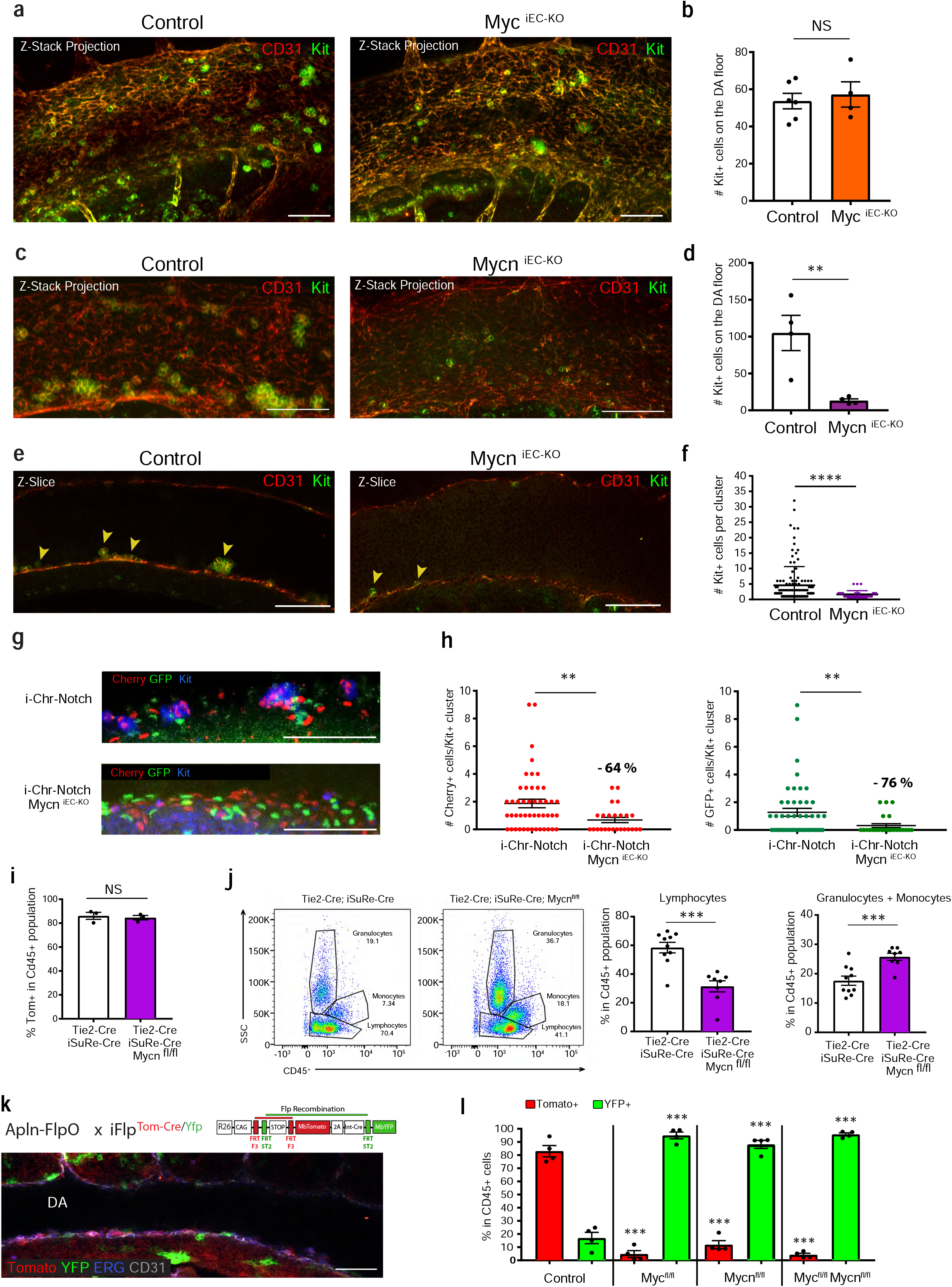
Mycn, unlike Myc, is required for EHT. a) Z-stack projection of confocal scans of control and *Myc*^*iEC-KO*^ E10.5 DAs, immunostained for CD31 and Kit. b) Myc loss does not alter the number of Kit+ cells in the DA floor (n=6 sibling control and n=4 mutant embryos).. c) Z-stack projection of confocal scans of control and *Mycn*^*iEC-KO*^ E10.5 DAs, immunostained for CD31 and Kit. d) Mycn loss significantly reduces the number of Kit+ cells in the DA floor (n=4 sibling control and mutant embryos). e) Z-confocal scanning slice of control and *Mycn*^*iEC-KO*^ E10.5 DAs, showing individual aortic hematopoietic (Kit+) clusters. f) Mycn loss significantly reduces the number of Kit+ cells per cluster (n=4 sibling control and mutant embryos). g) Z-stack projection of confocal scans of *iChr2-Notch-Mosaic* and *iChr2-Notch-Mosaic Mycn*^*iEC-KO*^ E10.5 DAs, immunostained for Cherry, GFP and Kit. h) Quantification of Cherry+ and GFP+ cells per Kit+ cluster, showing decreases after loss of *Mycn*, i) Frequency of Tomato+/iSure-Cre+ cells in the CD45+ population of adult animals with loss of Mycn driven by Tie2-Cre (targeting endothelial and hematopoietic cells), showing no significant difference from controls (n=3 sibling control and mutant mice). j) Mycn loss induces a decrease in the lymphocyte population and an increase in the granulocyte and monocyte populations, suggesting lineage bias (n=10 sibling control and n=8 mutant mice). k) Mice carrying the alleles *Apln-FlpO* and *iFlp*^*MTomato-Cre/MYfp*^ have induction of an EC mosaic in the DA at E10.5. Tomato+ cells express Cre and become mutant when in a floxed gene background, whereas YFP+ cells have no genetic deletion (internal control). l) Comparative analysis of the relative percentage of Tomato+ and YFP+ cells determined by FACS in the blood of adult mice carrying the *Apln-FlpO* and *iFlp*^*MTomato-Cre/MYfp*^ alleles alone (Control) or in combination with the indicated floxed alleles. Given the shorter genetic distance between the FRT sites present before the MbTomato-2A-Int-Cre cassette, about 80% of blood cells are Tomato+ on the non-floxed background (Control), whereas only about 20% are YFP+. When these alleles are on a *Myc*^*fl/fl*^ or *Mycn*^*fl/fl*^ background, only Tomato-2A-Cre+ cells have a permanently induced genetic deletion (see also Extended Data Fig. 1). Loss of *Myc* or *Mycn* in DAECs and their hematopoietic descendents (Tomato+) strongly impacts hematopoietic development (n=4 sibling control and mutant mice). Scale bars, 100 μm. Error bars indicate SEM. NS, non significant; **p < 0.01; ***p < 0.001; ****p < 0.0001.

Mycn is molecularly and functionally similar to Myc, completely rescuing its function when inserted in the *Myc* locus ^49^. Given the strong expression of *Mycn* during EHT (Fig. 6g, 6h), we analyzed *Mycn Tie2-Cre (Mycn*^*iEC-KO*^) E10.5 embryos. In contrast to single *Myc*^*iEC-KO*^ mutants, *Mycn*^*iEC-KO*^ embryos showed a significant reduction in the number of Kit^+^ HPCs in the DA endothelium (Fig. 7c, 7d). We also investigated HPC cluster growth dynamics and saw that besides *Mync*^*iEC-KO*^ embryos having fewer clusters, they were also significantly smaller (Fig. 7e, 7f). These results indicate that *Mycn*, not *Myc*, is a key driver of EHT.

We then wanted to determine if the *Mycn* upregulation seen in DA ECs with lower Notch signaling (Fig. 6e) is indeed a key event for these cells to undergo EHT. For that we crossed *Tie2-Cre* mice with *iChr2-Notch-Mosaic* and *Mycn* floxed mice. In this way we were able to quantify how the GFP^+^/DN-Maml1^+^ mutant aorta ECs differentiated and expanded in the absence of *Mycn*. The results show that the loss of *Mycn* induces a significant decrease in the *iChr2-Notch-Mosaic* cluster size, and particularly reduces the contribution of GFP^+^/DN-Maml1^+^ cells to the clusters (Fig. 7g, 7h). This shows that Mycn is indeed a critical regulator of EHT of DAECs with lower Notch signaling.

### Mycn and Myc functions are required sequentially during embryonic hematopoiesis

The strong EHT defect identified in *Mycn*^*iEC-KO*^ embryos suggested that *Mycn* deletion was incompatible with the development of the adult hematopoietic system. Embryos lacking the function of *Myc* in Tie2-Cre+ cells die at E12.5 and have marked vascular and hematopoietic development defects^51^. However, vascular or hematopoietic defects have not been reported in global *Mycn* knockout embryos, which die by E11.5 with development defects in the brain, heart, lung, and genitourinary system ^52, 53, 54, 55, 56, 57^. We found that, unlike *Myc*^*iEC-KO*^ mice, *Mycn*^*iEC-KO*^ adult mice were viable and formed blood. Given that some HPCs formed and expanded in *Mycn*^*iEC-KO*^ aortas (Fig. 7c, 7d), we first hypothesized that the survival of *Mycn*^*iEC-KO*^ adults and their lack of hematopoietic defects was due to incomplete expression of *Tie2-Cre* (as shown in Fig1e, 1f) and incomplete *Mycn* deletion in aortic ECs during EHT. To exclude this possibility that blood in *Mycn*^*iEC-KO*^ adults derives from wildtype DA EHT events, we crossed *Mycn*^*iEC-KO*^ mice with *iSuRe-Cre* mice, in which cells expressing MbTomato-2A-Int-Cre have full deletion of any floxed allele ^58^. Surprisingly, adult *Mycn*^*iEC-KO*^ *iSuRe-Cre* and control *Tie2-Cre iSuRe-Cre* mice had similar numbers of Tomato-2A-Cre+ hematopoietic cells (Fig. 7i). The only detected differences in *Mycn*^*iEC-KO*^ *iSuRe-Cre* mice were an almost 2-fold decrease in lymphocytes as a percentage of the total CD45^+^ population and a corresponding increase in the percentage of myeloid cells (Fig. 7j). These adult blood data are incompatible with our earlier embryo EHT studies, and we therefore used a public server (gexc.riken.jp) to check for the expression of *Tie2* (*Tek)*. This analysis revealed that *Tie2* is expressed not only in ECs, but also in the blood lineage, with particularly high expression in adult LT-HSCs. Therefore, the high percentage of Tomato^+^ cells in the CD45^+^ blood population of adult *Mycn*^*iEC-KO*^ mice might reflect recombination of the *iSuRe-Cre* reporter, with subsequent loss of *Mycn*, not in early hemogenic ECs but at later embryonic stages or in adult HSCs.

To assess the impact of *Mycn* loss-of-function on DAEC-derived adult blood, we induced genetic mosaics with two new mouse alleles (*Apln-FlpO* and *iFlp*^*Tom-Cre/Yfp*^*-Mosaic*) generated in our laboratory (Garcia-Gonzalez et al., in preparation). The combination of these 2 alleles with *Mycn-*floxed alleles allowed us to induce a trackable mosaic of control cells (YFP+) and *Nmyc*^*KO*^ cells (MbTomato-2A-Cre+) derived from Apln+ ECs. *Apln-FlpO* is an endogenous X-chromosome locus knock-in allele subject to mosaic inactivation in female embryos, and therefore expressed only in a fraction of embryonic Apln+ ECs, including those that will form the DA at E10.5 (Fig. 7k). Unlike *Tie2, Apln* is not expressed in subsequent hematopoietic progenitors or blood lineages (gexc.riken.jp). In control mice, with the *Apln-FlpO* and *iFlp*^*Tom-Cre/Yfp*^*-Mosaic* alleles, *Tomato-2A-Cre*^*+*^ DAECs contributed about 80% of the total recombined blood cells, whereas YFP^+^ cells contributed only about 20% (Fig. 7l, control bars), which is consistent with the shorter genetic distance between the FRTF3 sites (Fig. 7k and Extended Data Fig.1). This Tomato:YFP ratio was very consistent between animals (n=4 adults). Examination of the Tomato:YFP ratio in *Myc*^*fl/fl*^, *Mycn*^*fl/fl*^, and *Myc/Mycn*^*fl/fl*^ adult blood revealed a significantly smaller Tomato-2A-Cre^+^ population; instead of 80%, only 5-10% of the total recombined CD45^+^ cells were Tomato^+^ (Fig. 7l).

The sum of the results obtained with the Apln-FlpO and Tie2-Cre driver lines confirms that single deletion of *Mycn* or *Myc* strongly impairs hematopoietic development; however, *Mycn* (and Notch) function is only relevant during EHT, and not essential at later stages of embryonic hematopoietic development, in line with its non-requirement for adult hematopoiesis ^59^. This finding matches the transient nature of *Mycn* expression during EHT and the relatively low *Mycn* expression in more developed hematopoietic progenitors and lineages (Fig. 6g). In contrast to *Mycn, Myc* expression increases as hematopoietic lineages develop, and *Myc*^*iEC-KO*^ mice have defects in hematopoietic expansion and development (Fig. 7l), but not in EHT or DA HPC cluster formation (Fig. 7a, 7b). Together, these data indicate that while *Mycn* is mainly important for EHT, *Myc* acts subsequently during hematopoietic lineage development.

## Discussion

Effective modulation of endothelial-to-hematopoietic transition may enable the generation of HSCs from ECs in sufficient numbers for use in regenerative medicine. This could be particularly useful in blood cancer or other hematopoietic disorders caused by mutations in hematopoietic stem or progenitor cells. Unlike those cells, ECs are quiescent and therefore less prone to the accumulation of genetic mutations, making them a potential autologous source of hematopoietic cells.

In this study, we combined multispectral functional genetic mosaics with single cell RNAseq data analysis to define the role of Notch signaling in EHT at high cellular and molecular resolution. Our findings differ from earlier reports in zebrafish embryos ^14, 15^ and human pluripotent stem cells^21^ as well as initial findings in mice ^8, 16^. Analysis of endothelial Notch genetic mosaics allowed us to find that increased Notch activity in DAECs inhibits EHT, a finding in line with more recent studies using distinct experimental approaches ^34, 60, 61^. We show that single ECs with mildly decreased or elevated Notch signaling can both form the first embryonic arteries, but their subsequent contribution to Kit+ hematopoietic clusters is influenced by their Notch signaling status. We found that once specified, arterial ECs with lower Notch signaling have a higher propensity to undergo EHT. Analysis of scRNAseq data revealed that Gfi1+ hemogenic arterial ECs have a pulsed increase in the expression of the Notch glycosyltransferase Mfng, which decreases the signaling ability of Jag1 ligands from neighboring cells ^25, 39^. The upregulation of Mfng expression and its glycosyltransferase activity on Notch receptors, lowers the level of Jag1-Notch signaling in signal-receiving hemogenic arterial ECs in a cell-autonomous manner, resulting in the strong upregulation of *Mycn* expression in AGM arterial ECs undergoing hematopoietic transition (Fig. 8 model). Our data suggest that Jag1 and Dll4 ligands expressed on DAECs compete for binding and activation of the glycosylated Notch receptors expressed by the hemogenic endothelium, leading to mutually antagonistic roles given their distinct signaling abilities ^25, 39^. Distinct and Fringe-dependent signalling outputs and functions for Jag1 and Dll4 were also observed during angiogenesis ^25^, endocardial development ^62^ and pancreas development ^63^. However, we found that Mfng expression in DAECs promotes EHT by decreasing Jag1-Notch signaling, whereas during angiogenesis Mfng mainly induces Dll4-Notch activation ^25^. This difference correlates with the relatively higher expression of Jag1 ligands in quiescent AGM DAECs, and the higher and VEGF-dependent expression of Dll4 ligands during angiogenesis ^64^.

**Figure 8.**
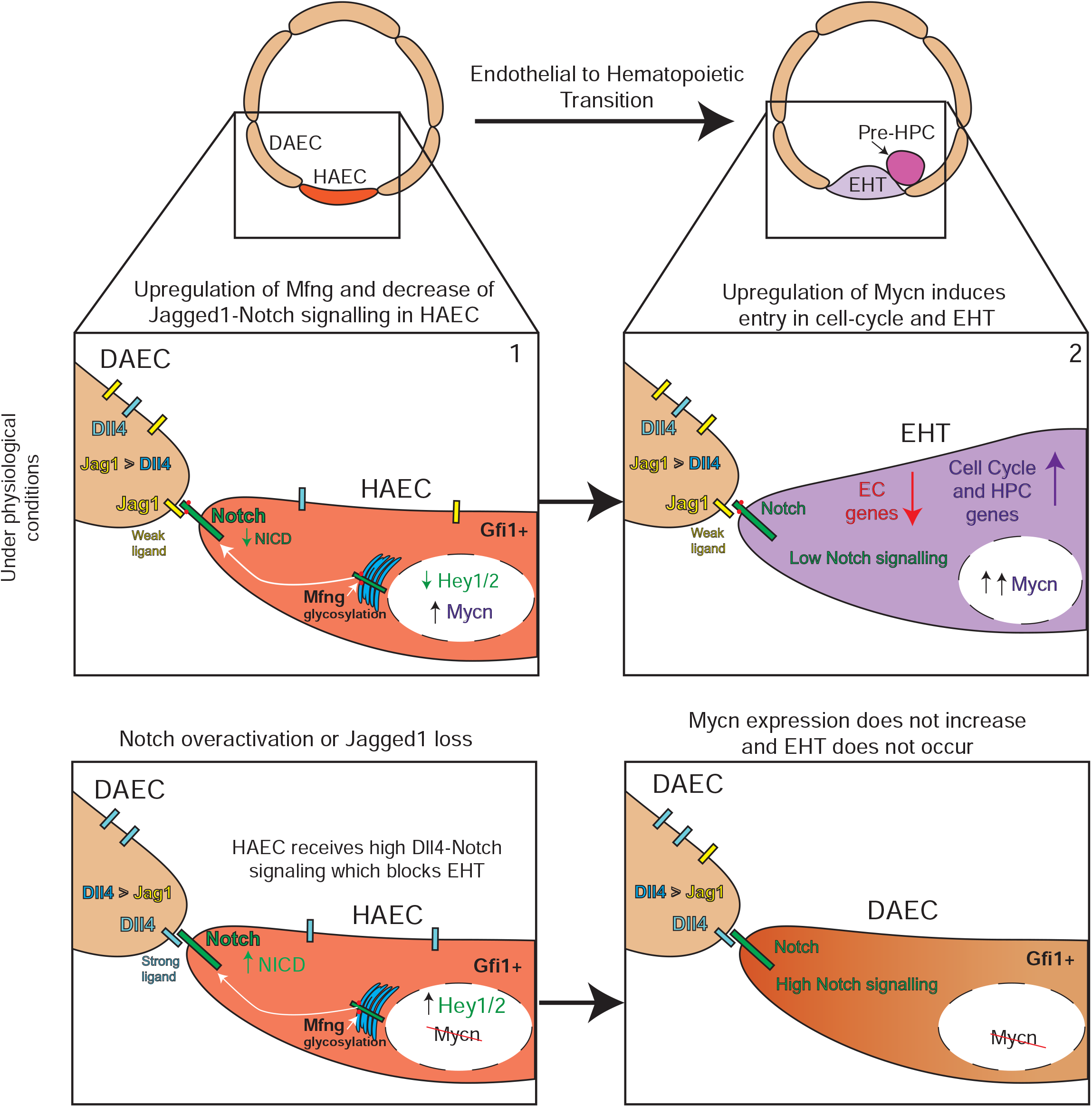
Model of EHT regulation by Notch and Mycn. A subset of dorsal aorta ECs located in the AGM region acquires hemogenic characteristics (Gfi1 expression) and has higher Mfng expression. Mfng glycosylates Notch receptors, rendering them refractory to signaling from Jag1 ligands expressed on neighboring AGM DAECs, which express Jag1 more abundantly than Dll4. The decrease in cell-autonomous Notch signaling activity in hemogenic arterial ECs (HAECs) leads to downregulation of Hey1 and Hey2 and upregulation of Mycn. As Mycn levels increase, the endothelial signature decreases and the cell-cycle and hematopoietic progenitor cell gene signatures increase, leading to endothelial-to-hematopoietic transition (EHT). Ectopic Notch activation or loss of Jag1 in DAECs increases Dll4–Notch signaling, resulting in higher Hey1 and Hey2 expression and the suppression of Mycn and EHT.

A previous single cell gene expression analysis by Baron and colleagues^40^ showed that *Mycn* is one of the most enriched genes in AGM cells undergoing EHT. Our transcriptional profiling of AGM DAECS with lower Notch signaling and a higher propensity to undergo EHT revealed the transcriptional upregulation of *Mycn* in these cells. Pseudotime analysis of the same single cell RNAseq dataset showed that this *Mycn* upregulation occurs right after the upregulation of *Mfng* and the downregulation of the Notch target gene *Hey2*, supporting the idea that hemogenic arterial cells with higher *Mfng* expression experience a decrease in Notch signaling before the upregulation of *Mycn*. This is the first report to show that Notch negatively regulates *Mycn* expression in AGM DAECs, contrasting the situation in hematopoietic cells, where Notch positively regulates the more widely known and homologous *Myc* gene ^48^. Importantly, we found that Mycn and Myc act sequentially during hematopoiesis. Our data indicate that Mycn is an important and specific regulator of EHT, whereas Myc plays a later role in hematopoietic lineage expansion. In contrast to Myc, Mycn function seems to be restricted to embryonic EHT and, like Notch signaling ^59, 65^, is not required for the homeostasis of the adult hematopoietic system. Within the endothelial lineage, Mycn function is also restricted to AGM ECs and EHT and is not an important regulator of other EC processes. This conclusion is supported by the finding that embryos with deletion of *Mycn* in *Tie2-Cre* lineages have no major developmental defects, unlike embryos with *Notch1* or *Myc* deletion ^18, 51^. Our competitive and long-term mosaic lineage tracing of DAECs with (YFP+) or without (Tomato and Cre+) Mycn function shows that cells lacking Mycn are strongly outcompeted due to the early impairment in EHT. Even though *Mycn Tie2-Cre* embryos eventually form HPCs and blood from the few aortic hematopoietic clusters that escape *Mycn* deletion, they have significantly more lymphocytes and fewer monocytes. Our results suggest that *Mycn* upregulation in single cells undergoing EHT promotes the early expansion of aortic hematopoietic clusters. Like Myc, Mycn promotes the expression of biosynthetic pathway genes that increase cell metabolism and cell proliferation, both processes being associated with EHT ^45^. The interconnection between cell proliferation and differentiation is common in developmental biology, and our data suggest that these processes are simultaneously triggered by the decrease in Notch signaling in the hemogenic endothelium.

It remains to be explored whether the simple act of inhibiting Notch signaling or forcing Mycn expression can induce any degree of EHT in other embryonic or adult arteries. Modulation of these and other mechanisms may one day enable the *in vivo* reprograming of adult ECs to HPCs. However, we first need to determine whether adult ECs have enough plasticity to be genetically reprogrammed and which combination of molecular signals would need to be activated in order to make them hemogenic.

**Extended Data Figure 1.**
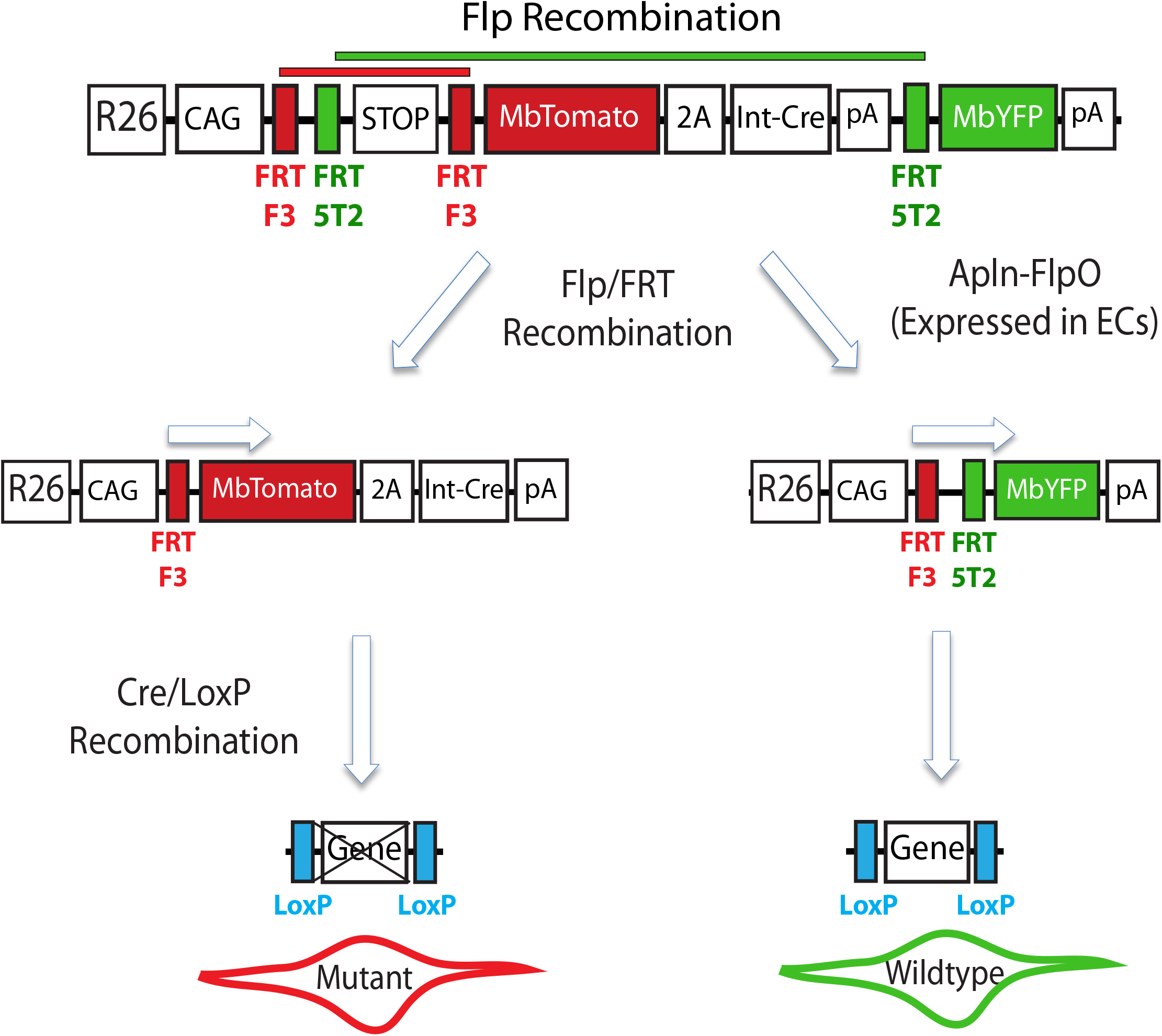
Inducing genetic mosaics with *iFlp*^*MTomato-Cre/MYfp*^ mice. Mice carrying the *iFlp*^*MTomato-Cre/MYfp*^ and *Apln-FlpO* alleles have induction of a genetic mosaic in Apln+ cells, such as endothelial cells. When cells express the recombinase FlpE or FlpO, a mutually exclusive recombination event occurs between the FRTF3 sites or the FRT5T2 sites. This generates a two-cell genetic-cellular mosaic. Some cells will express equimolar levels of MbTomato and Cre and the others will express only MbYFP. When these alleles are in a given floxed gene background, the MbTomato cells will have the gene deleted (KO) while the MbYFP cells will be wildtype.

## Materials and Methods

### Animals

*Gt(ROSA)26Sor*^*tm1(CAG-LSL-Tomato_GFP_N1ICDP)Ben*^ mice (abreviated *N1ICDP*^*iEC-GOF*^ when combined with *Tie2-Cre*) and the *Apln-FlpO and iFlp-Mosaic* mice (Luo et al., under review) were generated in our laboratory. We also used the previously published mouse lines *iChr2-Notch-Mosaic* ^31^; *Jag1*^*fl/fl* 66^; *Tie2-Cre* ^67^; *Rosa26-Lox-Stop-Lox-Mfng* ^62^; *Myc*^*fl/fl* 68^; *Mycn*^*fl/fl* 69^ and *iSuRe-Cre* ^58^. Embryos were obtained from timed pregnant females. The day of vaginal plugging was considered embryonic day (E) 0.5. Embryos were staged according to embryonic day, somite pairs (sp), and Thelier criteria (http://genex.hgu.mrc.ac.uk/intro.html). Mice and embryos were genotyped using the primers shown in Extended data table 1.

Animals were kept under pathogen-free conditions, and experiments were conducted in accordance with institutional guidelines and laws, following protocols approved by local animal ethics committees and authorities (Comunidad Autónoma de Madrid and UniversidadAutónoma de Madrid— CAM-PROEX 177/14 and CAM-PROEX 167/17).

### Tissue processing and whole-mount immunofluorescence

Embryos were transferred from the uterus to PBS, and the yolk sac was removed for genotyping. Embryos were staged according to the day of isolation or Thelier criteria. Dissected embryos were fixed in 2% PFA in PBS for 20 minutes at 4°C and then washed 3×15 minutes in PBS. Embryos were then dehydrated through increasing concentrations of methanol in PBS (50%, 75%, 100%; 30 minutes each) and stored in 100% methanol at −20C until processing. Embryos were further dissected as described ^70^. Briefly, the rostral half of the body, limb buds, and lateral body wall were removed in cold 100% methanol, and the dissected embryo tissue containing the AGM was rehydrated through decreasing methanol concentrations in ice-cold PBS (75%, 50%, 25%; 30 minutes each) following by 2 final washes inm ice-cold PBS. Tissues of mutant and control littermate embryos were blocked and permeabilized overnight at 4°C in blocking solution (1% skimmed milk, 0.4% Triton X-100, and 0.5% high-grade BSA in PBS). The day after tissues were incubated overnight at 4C with primary antibodies (Extended Data Table 2) diluted in PST (PBS, 1% skimmed milk, and 0.4% Triton X-100). After three 1 hour washes in PBT on ice, tissues were incubated overnight at 4°C in PBT containing Alexa-conjugated secondary antibodies (Extended Data Table 2). After three washes in PBS, embryos were dehydrated through a glycerol series (20%, 40%, and 60%) and cleared in cubic 1 solution^71^ for 24 hours. Embryos were then mounted in cubic 1: glycerol solution (1:1) in a FastWell chamber (FW20, Grace Bio-labs), coverslips were placed over the FastWells, and samples were stored at 4°C until imaging.

### Image acquisition

Up to 5 fluorophore signals per immunostaining were detected with a Leica SP8 confocal microscope, which is equipped with a white laser allowing excitation at any wavelength from 470 nm to 670 nm. For high-depth cleared AGM tissue imaging, we used excitation lines at 488 nm, 546 nm, 594 nm, 647 nm, and 680 nm and high-sensitivity hybrid detectors. In most cases, embryo tissues were immobilized in 1 mm deep FastWell chambers and were imaged from the lateral side. Images of the dorsal aorta (DA) were acquired to the left of the vitelline artery with a 20x lens and glycerol as the immersion medium. Confocal aquisition settings were a 2.5 Airy unit pinhole and a Z-step of 5.5-7.0 μM. All images shown are representative of results obtained in each group and experiment. Comparisons of phenotypes or immunosignals were made using images acquired with the same laser excitation and confocal scanner detection settings. The optimal Z-stack volume was determined for each embryo to ensure that it covered the entire depth of the dorsal aorta. Images were processed and quantified with the Fiji Image J-based image processing package.

### Flow cytometry and cell sorting

Flow cytometry analysis was performed in a FACS Aria Cell Sorter (BD Biosciences). For mRNA isolation from sorted cells, cells were incubated with either anti-CD31-APC (551262 BD Biosciences) or biotinylated anti-CD117/Kit^+^ (Tonobo Biosciences, 20-1172 1:300), and Rabbit-anti-mouse Connexin 40 (Alpha Diagnostic, CX40-A, 1:200) as primary antibodies, followed by Streptavidin-BV421-conjugated (BD Horizon, 563259 1:300) and Alexa Fluor 647-conjugated donkey anti-rabbit (Jackson Laboratories, 711-607-003 1:200) as secondary antibodies. ECs were sorted from the AGM and non-AGM CD31+ or Cx40^+^CD117^−^ populations and separated according to their endogenous fluorescence (GFP or Cherry). Sorted cells were collected in Qiagen RLT buffer, and mRNA was extracted using the Qiagen RNeasy Micro Kit (#74004).

For peripheral blood analysis, blood was drawn from the submaxilar region of mice older than 6 weeks and stained with anti-CD45.2-APC/CY7 (Tonobo Biosciences, 25-0454 1:100). Cells were analyzed or separated according to their endogenous fluorescence (GFP or Cherry).

### qRT-PCR

For quantitative real time PCR (qRT-PCR), RNA extracted from sorted ECs (see above), was retrotranscribed with the High Capacity cDNA Reverse Transcription Kit in the presence of RNase Inhibitor (Thermo Fisher, 4368814). cDNA was preamplified with Taqman PreAmp Master Mix containing a pool of Taqman Assays for the following genes: Actb, Gapdh, Hey1, Hey2, Efnb2, Cdh5, Myc, Mycn, Jag1 and Dll4 (see Extended Data Table 3, Thermo Fisher). Preamplified cDNA was used to perform a standard qRT-PCR with the same gene-specific Taqman Assays (Thermo Fisher) in an AB7900 thermocycler (Applied Biosystems).

For *i-Chr-Notch-Mosaic* EC profiling and comparison, RNA from GFP+ (DnMaml1+) cells was compared with RNA from Cherry+ (Control) cells. For the *N1ICDP*^*iEC-GOF*^ analysis, RNA levels in GFP+ cells was compared with the levels in GFP-cells from the same sorting session.

### Single cell transcriptomic analysis

The Seurat R package^72^ was used for clustering analysis and the Palantir python framework^73^ for pseudotime trajectory analysis. In brief, a matrix of counts from E10 developing aorta samples was downloaded from a CEL-Seq single cell experiment perform by Baron and colleagues ^40^(GSE112642). Paired end reads obtained by CEL-seq were aligned to the transcriptome using bwa (version 0.6.2-r126) with default parameters. The transcriptome contained all RefSeq gene models based on the mouse genome release mm10 downloaded from the UCSC genome browser and contained 31,109 isoforms derived from 23,480 gene loci. All isoforms of the same gene were merged to a single gene locus. The right mate of each read pair was mapped to the ensemble of all gene loci. Reads mapping to multiple loci were discarded. The left read contains the barcode information: the first eight bases correspond to a sample-specific barcode. The remainder of the left read contains a polyT stretch followed by few (<15 transcript) derived bases. Genes with at least 1 count in 6 cells and cells with 500 or more detected genes were kept for the analysis. Counts were normalized, log transformed and scaled. Groups of 546 cells were clustered with the graph-based algorithm provided by Seurat with the 10 first components of the PCA for the 715 most variable genes at a resolution of 1. The tSNE dimensionality reduction method was used at a perplexity setting of 100. The MAST method with the celular detection rate as covariate was used to detect markers for each cluster. Cellular type annotation was based on the original manuscript and manual inspection of markers.

Palantir-based pseudotime trajectory analysis was performed according to guidelines (https://github.com/dpeerlab/Palantir), starting with normalization and log transformation, followed by PCA and diffusion map calculation, and finally application of the Palantir algorithm, starting with the cell with highest expression of Vwf, an endothelial marker ^74^. For gene expression plots and trend expression across pseudotime, gene expression imputation was performed using MAGIC ^75^ based on the diffusion map and the Palantir-calculated pseudotime trajectory. The selected list of genes used for the cell-cycle (G1/S phase) analysis between clusters C0, C2, and C3 (Fig.6b) is taken from Torin et al., 2016.

### Statistical analysis

No randomization or blinding was used, and animals or tissues were selected for analysis based on their genotype and the quality of immunostaining and confocal images. Statistical analyses were performed in GraphPad Prism. Comparisons between two samples with a normal distribution were made by unpaired two-tailed Student *t-*test or, when the standard deviation varied, unpaired *t-*test with the Welch correction. Data are presented as means ± SEM, unless otherwise specified. Population differences were considered statistically significant at P <0.05. Sample size was chosen for each experiment according to the observed statistical variation.

## Author Contributions

B.L. and R.B. designed experiments and interpreted results. B.L. executed the vast majority of the experiments. I.G.G engineered the DNA constructs and validated the *iFlpMosaic* and *Apln-FlpO* mice. V.C.G. gave general technical assistance and established some of the ES cell lines used to generate mouse models. R.B. and B.L. wrote the manuscript.

## Competing Interests

The authors declare no competing interests.

## Acknowledgments

We thank Simon Bartlett for English editing; Sofia Sanchez for assistance with the mouse colony and genotyping; Patricia Sanchez for her preliminary findings with *N1ICDP*^*iEC-*GOF^ embryos; the CNIC Transgenesis Unit for support in the generation of the mouse lines; the CNIC Microscopy Unit for confocal microscopy imaging; the CNIC Bioinformatics Unit, particularly Carlos Torroja Fungairino, for the single cell transcriptomic analysis. Research was supported by the European Research Council (ERC-2014-StG-638028), the Centro Nacional de Investigaciones Cardiovasculares (CNIC), and the Ministerio de Economia, Industria y Competitividad (MEIC: SAF2013-44329-P, SAF2013-42359-ERC and RYC-2013-13209). The CNIC is supported by the Ministerio de Ciencia e Innovación (MCIN) and the Pro CNIC Foundation and is a Severo Ochoa Center of Excellence (SEV-2015-0505).

---

**Extended Data Table 1.**
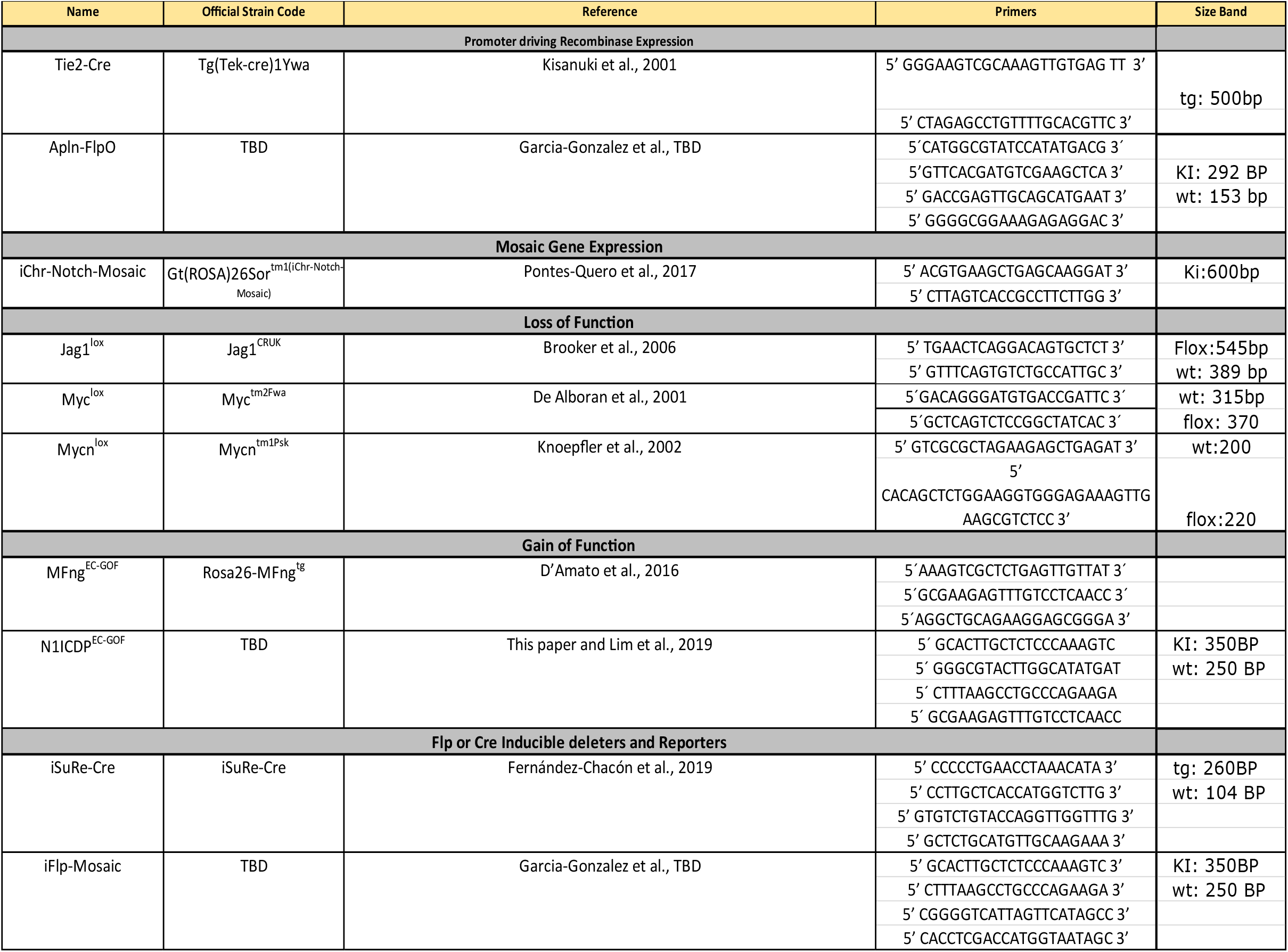
-Mouse Lines and Genotyping

**Extended Data Table 2.** -Antibody References

| Extended Data Table 2 - Antibody References |  |  |
| --- | --- | --- |
| Antibody | Company | Reference |
| *Primary Antibodies* |  |  |
| Goat anti-GFP | Acris Antibodies | Cat# R1091P |
| Rabbit anti-DsRed | Clontech | Cat# 632496 |
| Hamster anti-CD31 | Millipore | Cat# MAB1398Z |
| Mouse anti-HA-AF647 | Cell Signaling | Cat# 3444S |
| Rabbit-anti Connexin 40 | Alpha Diagnostic | Cat # 28cx4029 |
| Rabbit anti-RUNX1 | Abcam | Cat# ab92336 |
| Rabbit anti-Myc | Millipore | Cat# 06-340 |
| Goat anti-Dll4 | R&D | Cat# AF1389 |
| Rabbit anti-Kit | Abcam | Cat# ab16667 |
| Rabbit anti-ERG | Abcam | Cat# ab110639 |
| Rabbit anti-ERG-AF647 | Abcam | Cat# ab196149 |
| Rat anti-CD31-APC | BD Biosciences | Cat# 551262 |
| Rat anti-CD117/Kit-APC | Tonobo Biosciences | Cat# 20-1172 |
| CD45.2-APC-Cyanine 7 | Tonobo Biosciences | Cat# 25-0454-U025 |
| Cd11b-APC | BD Pharmigen | Cat# 51_01712J |
| B220-Biotin | BD Pharmigen | Cat# 51_0122J |
| CD3e-Biotin | BD Pharmigen | Cat# 51_01082J |
| *Secondary Antibodies* |  |  |
| Donkey anti-Goat AF488 | Thermo Fisher | Cat# A-11055 |
| Donkey anti-Rabbit AF594 | Thermo Fisher | Cat# A-21207 |
| Donkey anti-Rat AF647 | Abcam | Cat# ab150155 |
| Goat anti-Armenian Hamster<br>AF680 | Jackson Laboratories | Cat# 127-625-160 |
| Donkey anti-Rat CF633 | Biotium | Cat# 20137-1mg |
| Donkey anti-Rabbit CY3 | Jackson Laboratories | Cat# 711-167-003 |
| Streptavidin-BV421 | BD Horizon | Cat# 563259 |

**Extended Data Table 3.**
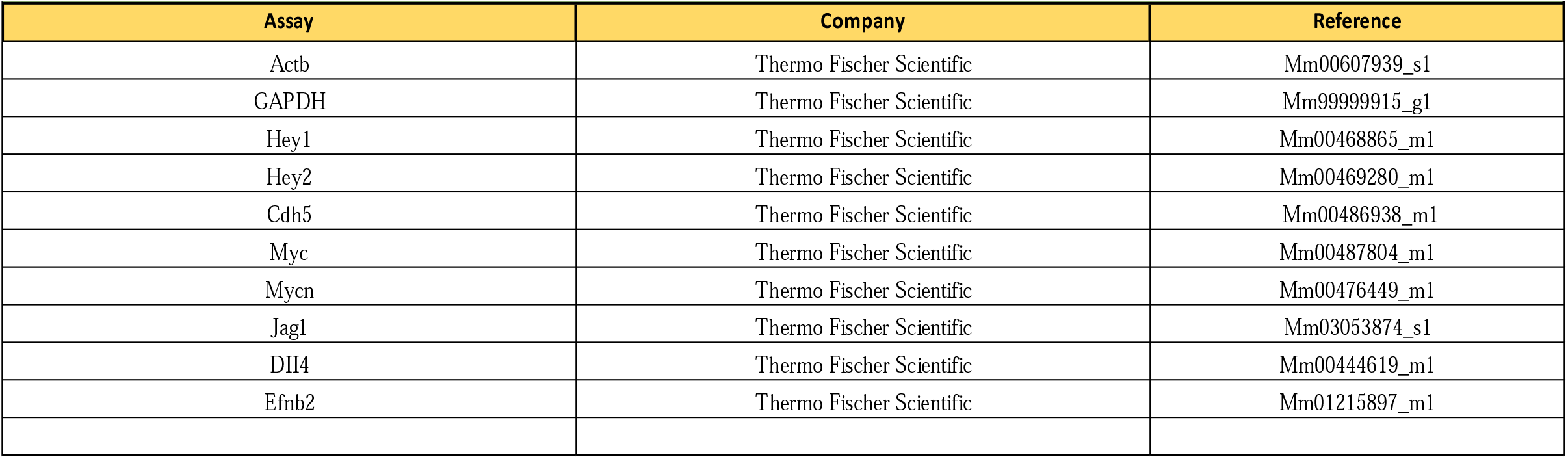
-TaqMan Assays/Probes

## Notes

### Competing Interest Statement

The authors have declared no competing interest.

